# Modulating CRISPR-Cas genome editing using guide-complementary DNA oligonucleotides

**DOI:** 10.1101/2022.01.15.475214

**Authors:** Thomas Swartjes, Peng Shang, Dennis van den Berg, Tim A. Künne, Niels Geijsen, Stan J.J. Brouns, John van der Oost, Raymond H.J. Staals, Richard A. Notebaart

## Abstract

CRISPR-Cas has revolutionized genome editing and has a great potential for applications, such as correcting human genetic disorders. To increase the safety of genome editing applications, CRISPR-Cas may benefit from strict control over Cas enzyme activity. Previously, anti-CRISPR proteins and designed oligonucleotides have been proposed to modulate CRISPR-Cas activity. Here we report on the potential of guide-complementary DNA oligonucleotides as controlled inhibitors of Cas9 ribonucleoprotein complexes. First, we show that DNA oligonucleotides down-regulate Cas9 activity in human cells, reducing both on and off-target cleavage. We then used *in vitro* assays to better understand how inhibition is achieved and under which conditions. Two factors were found to be important for robust inhibition: the length of the complementary region, and the presence of a PAM-loop on the inhibitor. We conclude that DNA oligonucleotides can be used to effectively inhibit Cas9 activity both *ex vivo* and *in vitro*.

## Introduction

CRISPR-Cas (clustered regularly interspaced short palindromic repeats and CRISPR-associated proteins) systems provide prokaryotes with adaptive immunity against mobile genetic elements (MGEs) ^1,2^. Similar to other CRISPR-Cas systems, the Cas9 nuclease from *Streptococcus pyogenes* (SpCas9) mediates double stranded cleavage of the target DNA ^3,4^. In brief, the Cas protein binds a guide RNA (gRNA) to form a ribonucleoprotein (RNP) complex. This complex then interrogates the DNA to find a DNA sequence complementary to the gRNA (protospacer) ^5^. To this end, the RNP complex binds DNA sequences with a protospacer adjacent motif (PAM) and causes initial unwinding of the adjacent DNA bases ^3–8^. If these bases are complementary to the crRNA, DNA unwinding proceeds along the gRNA until a stable R-loop is formed ^5,9^. Lastly, both DNA strands are cleaved, resulting in a double-strand break (DSB) in the target DNA ^3,4^.

Cas9 and other Cas nucleases are used for genome editing by introducing DSBs at specific DNA sequences. One possibility is that the genomic edits occur independently of the Cas nuclease. The nuclease would then be used as a counter-selection system to select against non-edited versions of the target site by introducing DSBs ^10^. Alternatively, the Cas nuclease might first introduce a DSB, which then induces local DNA repair ^11^. The two most common repair mechanisms that act on DSBs are homology directed repair (HDR) and non-homologous end joining (NHEJ) ^12^. HDR uses a homologous repair template to fix the DSB according to the template sequence ^12^. NHEJ resolves the double-stranded break without the need for a repair template, which often results in insertions or deletions (indels) at the site of the DSB formation ^13^.

For genome editing applications of CRISPR-Cas, it is crucial that the Cas nuclease introduces DSBs at the sequence of interest with sufficient efficacy. However, Cas-nucleases were found to also create DSBs at sequences with imperfect complementarity to the gRNA ^14–18^. It is essential to prevent genetic changes at such off-target sites, especially for therapeutic genome editing. In addition, strict control over Cas9 activity might be used to confine DSB formation to the desired cells in a limited time-frame.

In nature, Cas enzyme activity can be inhibited by phage-encoded anti-CRISPR (ACR) proteins that act on different stages of CRISPR-Cas based immunity ^19–21^. The ACRs that were found to inhibit Cas nuclease activity, have been reported to be useful for controlling CRISPR-Cas-based genome editing ^21–26^. Aside from these naturally occurring ACRs, several other strategies have been devised to control Cas enzyme activity at the level of transcription ^27–29^ , translation ^30–32^ , protein state ^33–47^ , and guide RNA ^14,48^.

In addition, single stranded DNA or RNA molecules can be designed to inhibit Cas nucleases. Such oligonucleotide-based inhibitors provide several advantages compared to natural ACR proteins. DNA oligos are inexpensive, can be rapidly manufactured, and could provide a systematic way to inhibit different Cas nucleases, whereas the use of anti-CRISPRs is dependent on compatibility with the Cas nuclease of choice.

Oligo-based inhibitors have been shown to work *in vitro* and in human cell cultures for Cas9 and Cas12a ^49–51^. Potent inhibition of Cas9 was observed with RNA-DNA hybrids or chemically-modified DNA inhibitors which interact with the repeat sequence of the guide RNA or with the PAM-interacting domain of Cas9 ^50^. In addition, truncated gRNA designs have been shown to allow dsDNA binding, but not cleavage by Cas9 ^52,53^. Such truncated gRNAs were used to specifically direct non-cleaving Cas9 RNPs to off-target sequences, thereby preventing active Cas9 RNP from binding these off-target sites ^51^. Lastly, DNA oligos with phosphorothioate linkages displayed strong inhibition of Cas12a activity, apparently independent of the nucleotide sequence of the inhibitor used ^49^.

In the current study, we assessed whether SpCas9 could be inhibited with guide-complementary DNA oligonucleotides without any chemical modifications. To this end, we designed and tested different oligo-based inhibitors complementary to the spacer-derived part of the guide RNA (these oligos thus have the same sequence as the PAM-proximal part of the protospacer). We also investigated the effect of extending the inhibitors with a double-stranded PAM sequence. We show that various designs provide strong, sequence-dependent inhibition of Cas9 *in vitro* and *ex vivo*.

In addition, we investigated the effect of oligo-based Cas9 inhibition on off-target sites in the context of genome editing. We found that while inhibition reduces both on- and off-target activity of Cas9, the presence of specific oligo designs results in slightly increased specificity. By comparing the inhibitor results with a Cas9 titration, we concluded that the increased specificity is a general consequence of lowering overall Cas9 activity.

Lastly, we studied which mechanisms lead to the observed inhibition. We concluded that the effect of the tested inhibitors is dependent both on the length of the oligo-based inhibitors and on the presence of a PAM-loop. Their relative importance is strongly affected by the speed at which Cas9 cleaves the targeted DNA.

## Results

To assess whether guide-complementary oligonucleotides could work as guide-specific inhibitors of Cas nucleases, we designed guide-complementary DNA oligos (hereafter referred to as ‘inhibitors’) of 8 and 20 nucleotides (nt) (Fig. 1). The inhibitors are complementary to the PAM-proximal, spacer-derived part of the guide RNA for SpCas9 (Fig. 1B,C). We also designed oligos with a 5’-extension intended to loop and fold back onto itself, creating a double-stranded PAM (Fig. 1D).

**Fig. 1.**
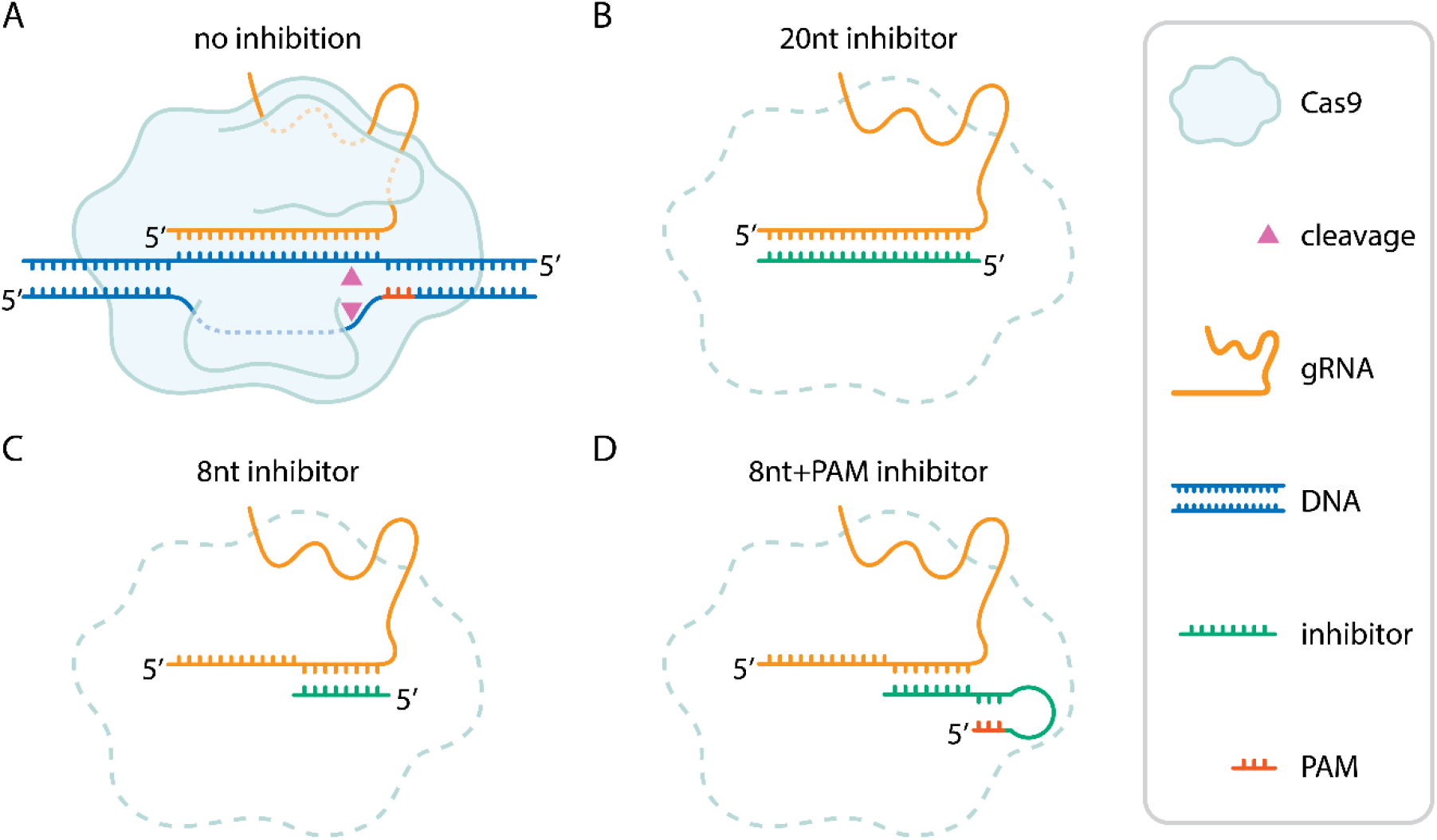
Design of guide-complementary oligo-based inhibitors. Schematic representation of a Cas9 nuclease, guide RNA (here displayed as a single guide RNA, sgRNA) with and without oligo-based inhibitors. (A) without inhibitors, the RNP binds the target sequence in genomic DNA, forms an R-loop and proceeds to cleave both strands of the DNA (triangles). (B) 20nt inhibitor. (C) 8nt inhibitor. (D) 8nt+PAM inhibitor. For all nucleic acids, 5’ ends are indicated.

We set out to test the effects of these DNA oligo-based inhibitors in a genome editing context. To that end, we delivered purified SpCas9 protein, guide RNA, and the inhibitors to Chronic myelogenous leukemia (CML) cells using iTOP ^54^. We analyzed editing of two endogenous gene loci, EMX1-1 and FANCF-2, under all test conditions. iTOP-transduced cells were allowed to grow to 90% of confluency, we then extracted the genomic DNA from the cells. From the genomic DNA, we amplified six regions of interest: the two on target sites (EMX1-1 and FANCF-2) and two, previously identified ^55,56^ , prominent off-target sites for each (Supplementary Table 1). We performed deep-sequencing on the amplicons to assess the frequency of indels arisen from NHEJ.

Without inhibitors, iTOP-delivered Cas9 RNP achieved roughly 40% and 70% on-target indel frequencies on EMX1-1 and FANCF-2 loci, respectively (Fig. 2A). Addition of the 20nt inhibitor - in equimolar ratio to the Cas9 RNP - strongly reduces indel frequency (to less than 5%), indicating that Cas9-induced DSB formation is repressed. Repression is also observed with the 8nt+PAM inhibitor, but to a lesser extent (to between 20% and 50%). The 8nt inhibitor however, unexpectedly showed increased indel frequencies rather than inhibition. The same was observed for most of the sequence-scrambled versions of the inhibitors, which were included as a control for sequence specificity. Together, this shows that both the 20nt and 8nt+PAM inhibitors repress Cas9 activity *ex vivo* in a sequence-specific manner.

**Fig. 2.**
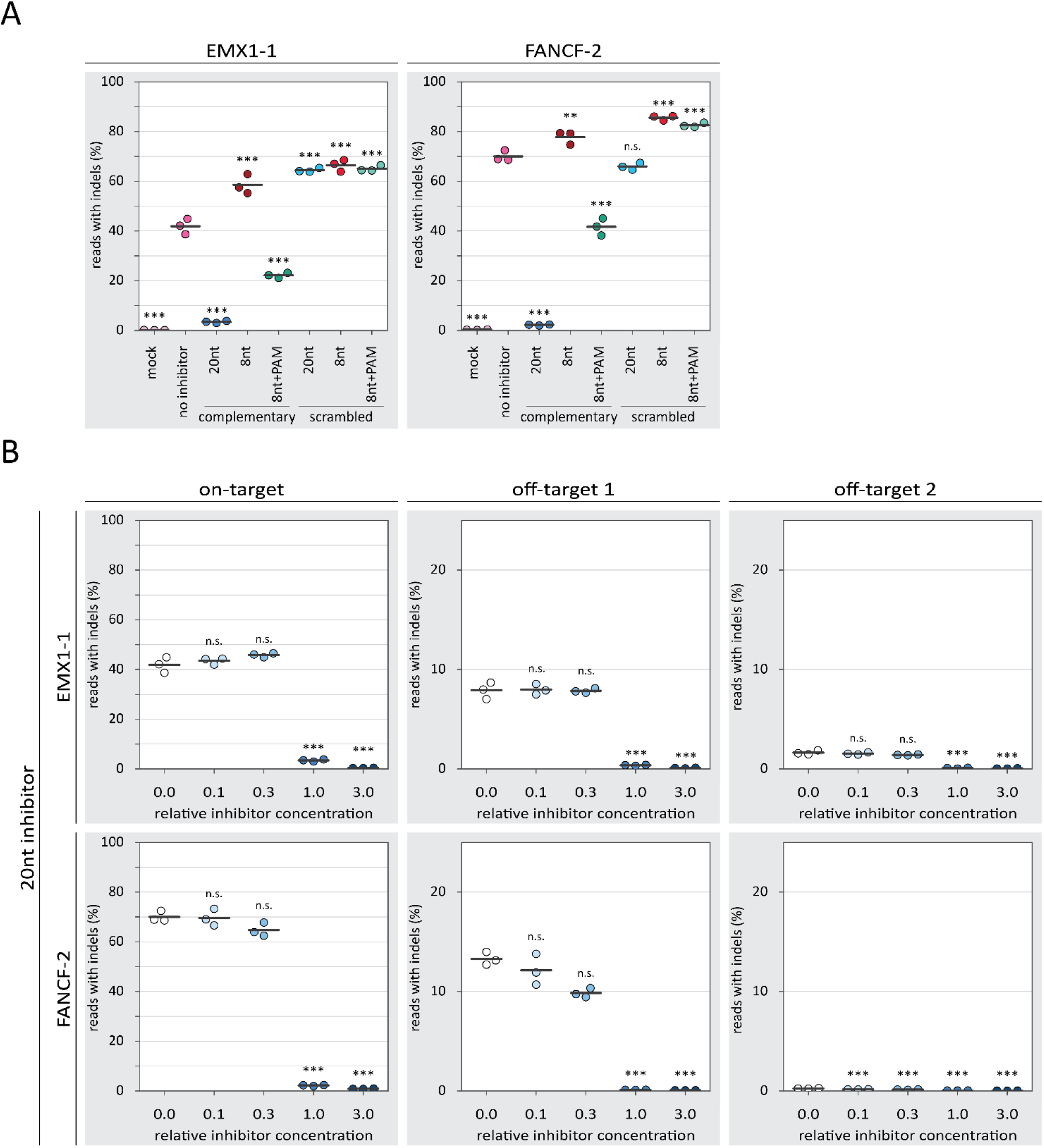
inhibition of Cas9 by guide-complementary DNA oligos. Individual replicates are displayed as colored dots while the horizontal black lines show the mean of the three replicates. *p*-values were calculated with a one-way ANOVA and subsequent Tukey’s test. ‘n.s.’: not significant; *: *p*-value < 0.05; **: *p*-value < 0.01; *** *p*-value < 0.001. **A:** Percentage of reads containing indels for on-target loci of EMX1-1 and FANCF-2. The ‘mock’ condition constitutes a control where the iTOP method was used without delivering Cas9, guide RNA and inhibitor. All conditions except the ‘mock’ used SpCas9, guide RNA and – except for ‘no inhibitor’ – a DNA oligo design as detailed in Fig. 1. The DNA oligos were delivered at a molar concentration equal to the concentration of Cas9 and guide RNA. *p*-values are displayed for the ‘mock’ condition compared to each other condition. **B:** Percentage of reads containing indels for on- and off-target loci of EMX1-1 and FANCF-2 for different concentrations of the 20nt inhibitor. The ‘relative inhibitor concentration’ indicates the molar concentration of the inhibitor relative to Cas9 and guide RNA. A ‘relative inhibitor concentration’ of 1.0 means that the used molar concentration of the inhibitor is equal to that of Cas9 and guide RNA. For off-target loci, the y-axis is adjusted because relatively few indels contained indels in these conditions. *p*-values are displayed for the 0.0 relative oligo concentration compared to each other condition.

To characterize the observed inhibition in more detail, we tested the inhibitors in a concentration gradient. Substantial inhibition by the 20nt inhibitor is only observed at and above equimolar ratio to the Cas9 RNP (Fig. 2B). For the 8nt+PAM inhibitors, a more gradual dose-response is observed (Fig. 3A) with low concentrations seemingly providing low levels of inhibition. However, even at the highest concentration, the 8nt+PAM inhibitors are not able to reduce indel frequencies to the same extent as was achieved with the 20nt inhibitors. At the investigated off-target loci, we observed reduced indel frequencies with the 20nt and 8nt+PAM inhibitors, following a similar dose-response as was seen on-target.

**Fig. 3.**
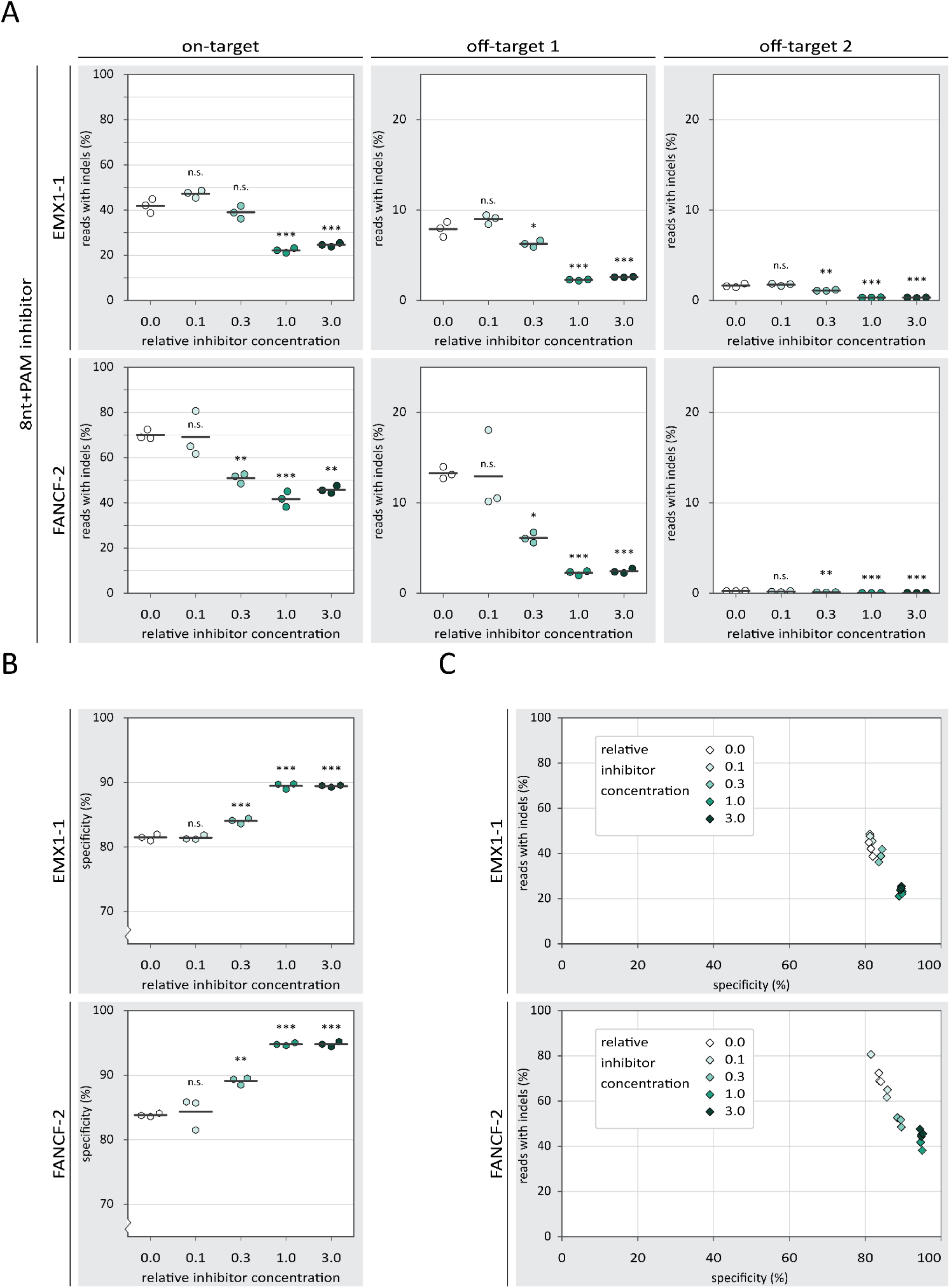
The effects of 8nt+PAM inhibitor concentration on specificity of Cas9-mediated NHEJ. *p*-values were calculated with a one-way ANOVA and subsequent Tukey’s test. *p*-values are displayed for the 0.0 relative oligo concentration compared to each other condition. ‘n.s.’: not significant; *: *p*-value < 0.05; **: *p*-value < 0.01; *** *p*-value < 0.001.**A:** Percentage of reads containing indels for on- and off-target loci of EMX1-1 and FANCF-2 for different concentrations of the 8nt+PAM inhibitor. The ‘relative inhibitor concentration’ indicates the molar concentration of the inhibitor relative to Cas9 and guide RNA. A ‘relative inhibitor concentration’ of 1.0 means that the used molar concentration of the inhibitor is equal to that of Cas9 and guide RNA. For off-target loci, the y-axis is adjusted because relatively few indels contained indels in these conditions. **B:** Percentage specificity for on-target loci of EMX1-1 and FANCF-2 for different concentrations of the 8nt+PAM inhibitor. The y-axis starts at 70% specificity to better visualize the differences between conditions. The individual replicates are displayed as colored hexagons while the horizontal black lines show the mean of the three replicates. **C:** comparison of *percentage reads with indels* and *percentage specificity* for different concentrations of the 8nt+PAM inhibitor. For each inhibitor concentration, individual replicates are displayed as diamonds with the same color.

The 8nt+PAM oligo displays inhibition both on- and off-target, but neither is completely abolished. This raises the question to what extent the ratio between on- and off-targeting is affected by the inhibitors. We calculated a specificity score by dividing the on-target indel frequency by the sum of the on- and off-target indel frequencies. This specificity score signifies the percentage of indels that were mapped to the on-target locus, corrected for the total number of reads mapped to each locus. We found that the higher concentrations of the 8nt+PAM oligo slightly increase the specificity score (Fig. 3B). However, it is important to realize that at the same inhibitor concentrations, on-target activity is substantially reduced (Fig. 3B,C).

We next asked whether the observed increase in specificity may be the result of a decreased subpopulation of active Cas9 RNP complexes through inhibition with DNA oligos. To address this, we looked at a range of Cas9 RNP concentrations to see how they affect on-target activity and specificity. At both on-target loci, we found that increasing RNP concentrations lead to higher indel frequencies (Fig. 4A). Strikingly, at the off-target sites, the highest Cas9 RNP concentration yielded relatively low indel frequencies. The resulting specificity scores are highest at relatively low RNP concentrations, and drop to a minimum at high concentrations. At the highest concentration however, specificity increases again, due to the unexpected decrease in off-target indels at that RNP concentration (Fig. 4B). When plotting both the on-target activity and specificity, among the intermediate Cas9 concentrations we observed a similar trade-off between activity and specificity as was observed with the 8nt+PAM inhibitor (Fig. 4C). The lowest and highest Cas9 RNP concentration do not follow this pattern however.

**Fig. 4.**
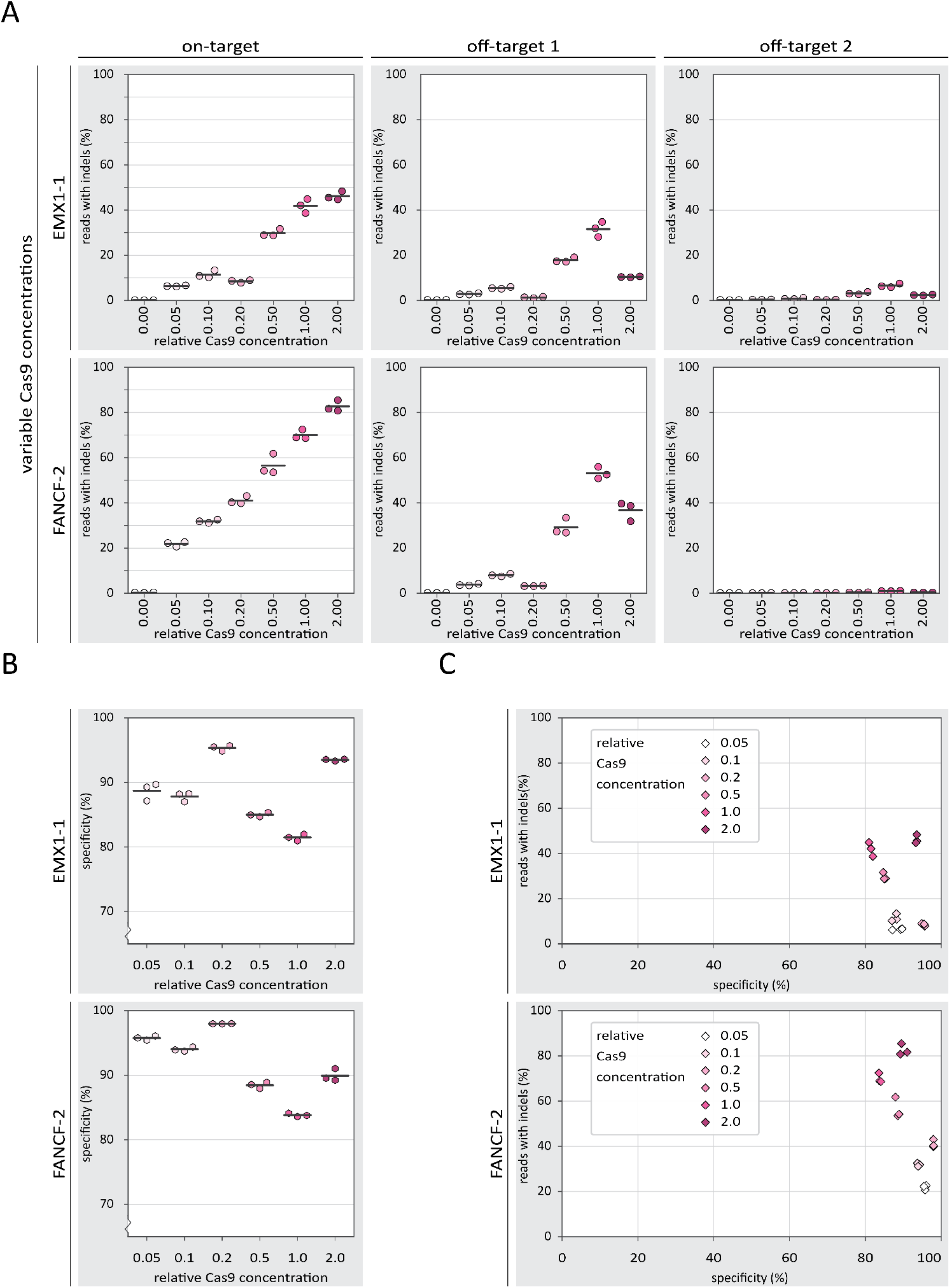
The effects of Cas9 concentration on specificity of Cas9-mediated NHEJ. **A:** Percentage of reads containing indels for on- and off-target loci of EMX1-1 and FANCF-2 for different concentrations of SpCas9 RNP in the absence of DNA oligo-based inhibitors. The individual replicates are displayed as colored dots while the horizontal black lines show the mean of the three replicates. The concentration 0.00 is the ‘mock’ condition in Fig. 2A. **B:** Percentage specificity for on-target loci of EMX1-1 and FANCF-2 for different concentrations of Cas9 RNP. The y-axis starts at 70% specificity to better visualize the differences between conditions. The individual replicates are displayed as colored hexagons while the horizontal black lines show the mean of the three replicates. The 0.00 Cas9 RNP concentration is not included here because the specificity values make no sense if there are virtually no indels at all. **C:** comparison of *percentage reads with indels* and *percentage specificity* for different concentrations of the Cas9 RNP. For each inhibitor concentration, individual replicates are displayed as diamonds with the same color. Again, the 0.00 Cas9 RNP concentration is not included here.

So far, we have seen that the 20nt inhibitor provides the most potent inhibition in our *ex vivo* setup, almost abolishing on- and off-target indel formation. The 8nt+PAM version displayed less inhibition, but shows some level of inhibition already at low concentrations. Next, we designed an experimental approach to better understand the mechanism by which the inhibitors function and shed light on why the 20nt and 8nt+PAM inhibitors display different dose-response *ex vivo*. To investigate the underlying mechanism, we used *in vitro* cleavage assays, which provide more experimental control than the *ex vivo* setup.

In preparation for the *in vitro* experiments, we performed PCRs of the EMX1-1 and FANCF-2 loci to produce the cleavage substrate DNA (supplementary table 4). Firstly, we incubated the purified Cas9 protein with the appropriate guide RNA (20 minutes pre-incubation at 25°C: Cas9 + gRNA). Secondly, we added the target dsDNA and the oligo-based inhibitors to the Cas9 and guide and incubated to allow cleavage of the DNA (30 minutes 37°C). We then used agarose gel electrophoresis to assess how much of the substrate DNA had been cleaved. We found that the pre-incubated Cas9 and guide RNA could efficiently digest the substrate DNA (Fig. 5). However, addition of the 8nt+PAM inhibitor substantially reduced DNA cleavage. In agreement with the aforementioned *ex vivo* analyses, we did not observe an effect from the 8nt inhibitor, nor with the sequence scrambled inhibitors. Unexpectedly, the 20nt inhibitor also did not show a clear reduction in DNA cleavage, despite strongly inhibiting indel formation *ex vivo*.

**Fig. 5.**
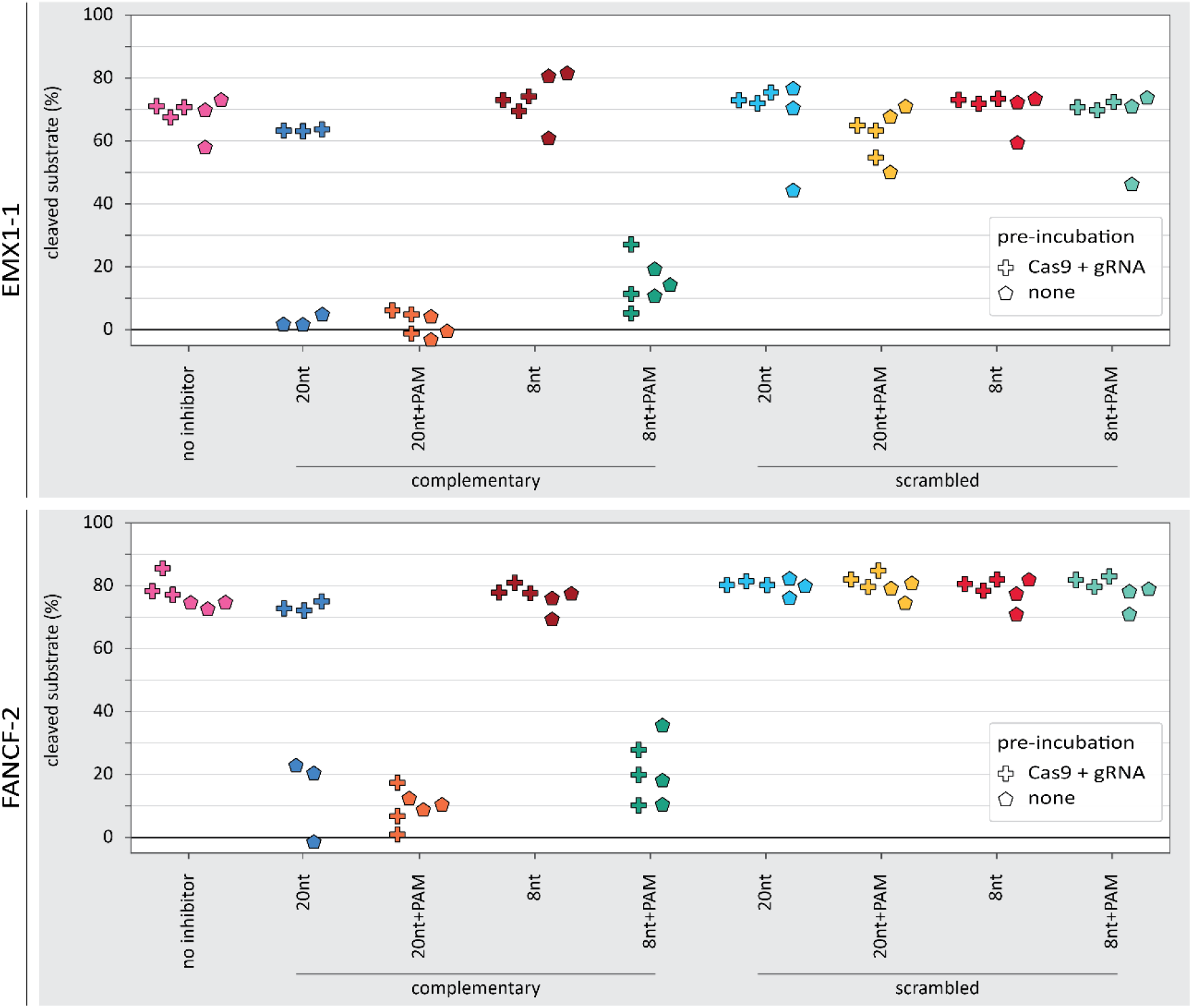
*in vitro* DNA cleavage in the presence of various oligo-based inhibitor designs. The percentage of DNA substrate cleaved as measured from band intensity on agarose gel. The plus-markers show the observed percentage of DNA cleavage when Cas9 and guide RNA were pre-incubated. The pentagon-markers represent replicates without pre-incubation. For EMX1-1 with oligo & gRNA pre-incubation, the scrambled oligo design each have a single datapoint that displays relatively poor cleavage. These low datapoints are all derived from a single batch of substrate DNA purified from agarose gel, different from the other datapoints.

To better understand the differences between *ex vivo* and *in vitro* outcomes we repeated the *in vitro* cleavage assays, but this time without the initial incubation of Cas9 with guide RNA (pre-incubation: none). In addition, we now included a 20nt+PAM inhibitor design in both experimental setups to see whether the PAM loop could rescue the performance of the 20nt inhibitor.

We now found that the 20nt inhibitor strongly inhibited DNA cleavage (Fig. 5). The 20nt+PAM inhibitor displayed strong inhibition in both experimental setups. This shows that inhibition without a PAM-loop is dependent on the order in which inhibitor, Cas9 and guide RNA are mixed. In contrast, the PAM-loop enables inhibition regardless of the mixing order.

We wondered whether the PAM-less 20nt inhibitor would only be able to bind the guide RNA when the RNA is not yet bound by Cas9. To assess this, we performed an electrophoretic mobility shift assay (EMSA) to visualize binding between inhibitor, Cas9 and guide RNA in different combinations. From this, we observed a large shift upon adding Cas9 to the guide RNA, indicating that the two bind to each other (Fig. 6A). If the 20nt inhibitor is added as well, the bands shift up slightly more, regardless of the order in which the components are mixed. This is not the case with the scrambled 20nt inhibitors, indicating that binding of the inhibitor is sequence-specific. Overall, we conclude that the 20nt inhibitor can bind both pre-formed RNP complexes and free guide RNA. It is therefore likely that the 20nt inhibitor can also inhibit at both stages.

**Fig. 6.**
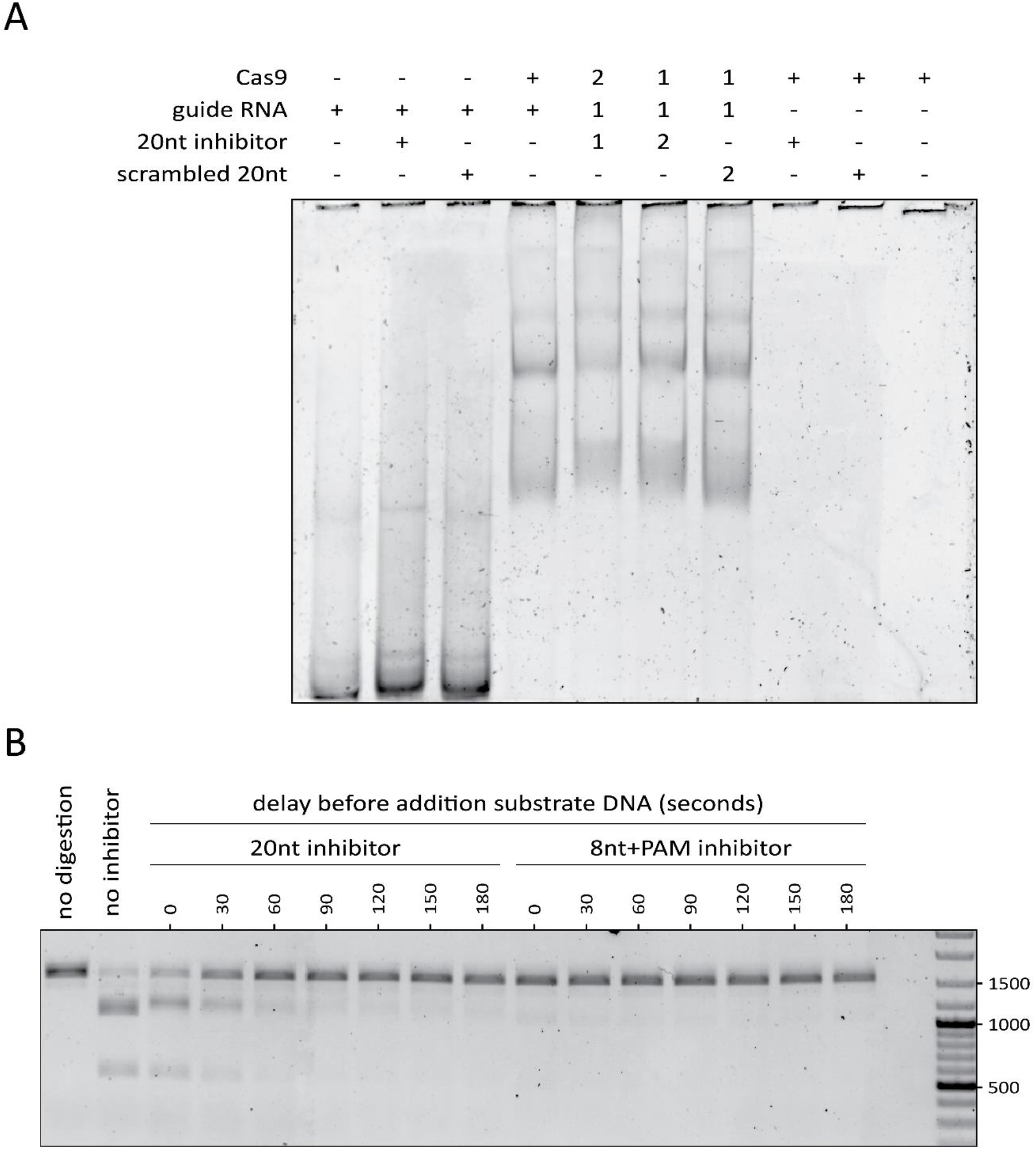
Timing-dependent inhibition of Cas9 by oligo-based inhibitors. **A:** electrophoretic mobility shift assay on 5% polyacrylamide gel, stained with SYBR Gold for nucleic acids visualization. Components with a ‘+’ sign were present, whereas those with a ‘-’ sign were not. A ‘1’ indicates that these components were initially mixed. In contrast, the components marked with ‘2’ were added after 15 min of initial incubation. The black lines at the top of the image correspond to the bottom of the wells of the gel. **B:** time-lapse assay where the 20nt or 8nt+PAM inhibitors were added to pre-formed RNP complexes, then incubated for variable durations (’delays’) before addition of the substrate DNA. The Generuler mix ladder was included on the far right and relevant fragment lengths are indicated in base-pairs. The linear substrate DNA is 1500bp long and cleavage by Cas9 would result in 2 fragments of lengths 1000bp and 500bp.

We then asked why – given the EMSA results – we did not observe inhibition with the 20nt inhibitor on pre-incubated Cas9 and guide RNA (Fig. 5). We reasoned that the 20nt inhibitor might require a longer time to bind pre-formed RNP compared to free guide RNA. If that is the case, the 20nt inhibitor would simply not have had enough time to bind (thus inhibiting cleavage of the DNA substrate by Cas9 RNP). We then tested how much time the 20nt inhibitor requires for inhibition of pre-formed Cas9 RNP. To that end, we first incubated the guide RNA and Cas9 to form the RNP. To this pre-formed RNP, we then incubated the 20nt or 8nt+PAM oligo for a variable duration before adding the substrate DNA and allowing 30 minutes of digestion.

We observed that the 20nt inhibitor requires roughly one minute to prevent most of the DNA cleavage upon subsequent addition of the substrate DNA (Fig. 6B). Inhibition by the 8nt+PAM design appears to be instantaneous. This explains why we did not observe a substantial reduction in DNA cleavage when the 20nt inhibitor was added simultaneously with the substrate DNA. Taken together, the *in vitro* results presented here show that a PAM-loop affects how the oligo-based inhibitors function. Designs with a PAM-loop can efficiently inhibit DNA cleavage regardless of whether they are added to free guide RNA, or to pre-formed RNP complexes. In contrast, PAM-less designs are effective when added to free guide RNA, and require a longer incubation to inhibit pre-formed RNP complexes.

## Discussion & Conclusion

We have described sequence-dependent inhibition of Cas9 with guide-complementary unmodified DNA oligos. In CML cells, we observed inhibition of on- and off-target NHEJ. Depending on the inhibitor design used, indels could be virtually eliminated. Additionally, we used *in vitro* assays to show that a PAM-loop enables robust inhibition by decreasing the time required for inhibition to take place. This PAM-loop design was inspired by another study where DNA oligos were used to create a double-stranded PAM ^57^. In contrast to most previously described Cas-inhibiting oligos ^49,50^ , the inhibitors described here inhibit in a guide sequence-dependent manner. Because of their sequence-specificity, these inhibitors theoretically allow modulating Cas9 activity at multiple sites independently. We found that the sequence-scrambled versions of the oligos (as well as the short (8 nt) complementary oligos) increased Cas9 activity, rather than inhibiting it. Perhaps the DNA oligos bind intracellular RNAs, preventing them from inhibiting RNP formation ^58^.

We found that the 20nt oligo is the most potent inhibitor tested in CML cells. The 20nt oligo outperforms shorter inhibitors, most likely because it has higher affinity to the guide RNA. The PAM-loop is thought to increase affinity, partially rescuing inhibition by shorter oligos. Unexpectedly, the 8nt+PAM consistently showed a slightly different concentration response compared to the 20nt inhibitor (Fig. 2B, 3A). This might be partially explained by assuming that the 20nt inhibitor is cleaved by Cas9, while the 8nt+PAM is not. Indeed *in vitro* ssDNA cleavage by SpCas9 has been described ^3^ and DNA cleavage requires interactions between the protein and ‘PAM distal’ bases of the DNA ^59^ , which are not present in the 8nt or 8nt + PAM inhibitors. This might allow the 8nt+PAM inhibitor to evade cleavage and inhibit Cas9 near-optimally at an equimolar ratio to the RNP. In cases where inhibition should last, a shorter-than 20nt oligo (perhaps with a PAM-loop to improve affinity) might therefore be preferred. Alternatively, chemical modifications - already proven to yield potent Cas nuclease inhibitors ^49,50^ - can prevent cleavage of inhibitors.

*In vitro*, the 20nt inhibitor design did not show efficient inhibition when RNPs were pre-complexed. Indeed, the PAM-less inhibitors required more time to inhibit pre-formed RNP compared to the designs with a PAM-loop. *Ex vivo*, the Cas9 protein and guide RNA likely entered the cell as free protein and RNA. Therefore, additional time was required to form RNPs, possibly providing the 20nt inhibitor more time for strong inhibition. In addition, in the CML cells, the on-target DNA substrate is presumably at a relatively low concentration compared to the in the *in vitro* conditions and – once the RNP is formed - the time required for the Cas9 RNP to find its cognate target is much longer. This too would give more time to the 20nt design to establish strong inhibition, possibly explaining the observed differences between the *ex vivo* and *in vitro* experiments.

We found that the DNA oligo-based inhibitors affect both on- and off-targeting by Cas9 and can slightly increase specificity for activity on-target. However, the observed increase in specificity is minor and coincides with a substantial reduction of on-target activity. To put these results into context, we also tested engineered Cas9 variants (SpCas9-HF1 ^60^ or Opti-SpCas9 ^61^) without inhibitors. We found that these variant Cas9 proteins are mostly more effective at increasing specificity than the inhibitors reported here (Supplementary Fig. 3). We thus conclude that for increasing specificity of Cas9, other measures (such as using engineered Cas9 protein variants) might be more suitable.

In an attempt to reveal the molecular basis of the specificity increase, we included conditions without inhibitors present where we varied the concentration of Cas9. The resulting data suggest that the observed increase in specificity could be a general consequence of lowering the concentration of active (cleavage-competent) RNPs, which has previously been described and explained by others ^15,62,63^. Unexpectedly, our highest Cas9 concentration led to increased on-target activity, but reduced off-target activity (Fig. 4A). The mechanism behind this is unknown and requires further research.

In CML cells, the most potent inhibition was observed with inhibitors that have the longest complementarity to the guide RNA. The *in vitro* experiments showed that a PAM-loop is required for rapid inhibition. Although the inhibitors performed slightly differently *in vitro* compared to *ex vivo*, in both settings DNA oligo-based inhibitors provided potent inhibition of Cas9 activity. Unmodified DNA oligos are inexpensive and easily manufactured and their design could easily be adapted to other RNA-guided nucleases. Overall, it is concluded that the guide-complementary DNA oligos reported here are promising candidates for inhibition of Cas9 activity.

## materials and methods

### Cell line and cell culture

The CML cell line used is a re-diploidized derivative of the HAP1 cell line, which was a kind gift of Dr. Thijn Brummelkamp ^64^. CML cells were cultured in Iscove’s Modified Dulbecco’s Medium (IMDM) (Gibco), supplemented with 10% fetal bovine serum and 1% penicillin/streptomycin. Cells were grown at 37°C in a humidified atmosphere containing 5% CO2.

### Cas9 protein, guide RNAs, and inhibitors used *ex vivo*

Recombinant *S. pyogenes* Cas9 and high-fidelity protein variants were provided by Geijsen lab through Divvly (https://divvly.com/geijsenlab). guide RNAs used are synthetic guide RNAs which contain the target-specific crRNA and the scaffold tracrRNA (IDT). The crRNA and tracrRNA were dissolved in the Nuclease-free Duplex buffer (IDT) to reach the concentration of 200 μM. Equal volumes of dissolved crRNA and tracrRNA were mixed and annealed by heating for 5 minutes at 95°C and cooling down at room temperature. The oligo-based inhibitors used were unmodified single-stranded DNA oligos synthesized by IDT (IDT). Each of such oligos was dissolved in nuclease-free water to reach the concentration of 75 μM.

### Induced transduction by osmocytosis and propanebetaine (iTOP)

The recombinant Cas9 proteins, guide RNAs, and oligo-based inhibitors were simultaneously transduced into CML cells by using the iTOP method as described previously ^54^. One day prior to the transduction, CML cells were plated at 18000 cells/well in the Matrigel-coated wells on 96-well plates, such that on the day of transduction, cells would reach about 70-80% confluence. Next day, for each well of the 96-well plate, 50 μL of iTOP mixture that contains 20 μL of transduction supplement (Opti-MEM media supplemented with 542 mM NaCl, 333 mM GABA, 1.67 x N2, 1.67 x B27, 1.67 x non-essential amino acids, 3.3 mM Glutamine, 167 ng/mL bFGF2, and 84 ng/mL EGF), 10 μL of Cas9 protein (75 μM), 7.5 μL of guide RNA (100 μM), 10 μL of oligo-based inhibitors, and the excess volume of nuclease-free water to reach the 50-μL total volume, were prepared. For the no-protein control, 10 μL of protein storage buffer was used instead of the CRISPR nuclease protein; and for the no-guide control, the equal volume of nuclease-free water was used to replace the guide RNA or oligo-based inhibitors. The 50-μL iTOP mixture was added onto the cells immediately after the culture medium was removed. The plate then was incubated in a cell culture incubator for 45 minutes, after which the iTOP mixture was gently removed and exchanged for 250 μL of regular culture medium.

### Isolation of genomic DNA in 96-well cell culture plates

Genomic DNA of each transduced cell samples was purified by direct in-plate cell lysis and DNA isolation according to a previously published protocol ^65^. Briefly, after aspirating culture media and adding 50 μL of lysis buffer (10 mM Tris HCl pH 7.6, 10 mM EDTA, 100 mM NaCl, 0.5% N-Lauroylsarcosine sodium, 50 μg/mL RNase A and 100 μg/mL Protease K) to each well, the sealed plate was incubated at 55°C overnight; added 100 μL of ice-cold NaCl-saturated ethanol (for 100 mL 100% ethanol, add 1.5 mL of 5M NaCl) to each well; allowed the plate stand still for 4 hours at room temperature to precipitate the DNA; washed the precipitated DNA with 75% ethanol for two times and let the plate air dry; DNA in each well was dissolved in 50 μL TE buffer.

### Deep-sequencing preparations

The regions of interest (supplementary Table 1) were PCR amplified (supplementary table 3) from the extracted DNA using Q5 high-fidelity DNA polymerase (NEB). Of the primers used in the PCRs (supplementary table 2) the forward primers contained 5nt sequencing barcodes. All 576 amplifications were verified using 20g/L agarose gel electrophoresis. Equal volumes of the PCR products were pooled with samples from the same locus, but with unique barcodes. These 36 pools were then purified (Zymo Research Z4004) and quantified using Qubit dsDNA BR (Thermo Fisher Scientific). Equal amounts of DNA from the samples were then pooled further to provide 6 pools where each sample contains a unique combination of sequencing barcode and amplified region. These samples were then sent to Baseclear B.V. for quality control, Index PCR, and sequencing on the NovaSeq 6000 for paired-end 150nt-long reads. The raw sequencing results are available on the NCBI Sequence Read Archive under BioProject PRJNA796802.

### Analysis of deep-sequencing results

To analyze the deep-sequencing data the paired-end reads were programmatically merged using *seqprep* ^66^. Then the reads were filtered out that do not match the expected pattern of starting with a barcode followed by a forward primer annealing part, and ending with the associated reverse primer annealing part. We then split the reads by their barcodes and mapped the reads to the human genome (hg38) using *bowtie2* ^67,68^. The resulting alignments were then – using *samtools* ^69,70^ - sorted, indexed, and split by the locus of interest where they align to the human genome. A separate script was used to score the amount of reads with insertions or deletions based on the CIGAR-strings from the alignment files. Another script was used to – for each indel – record its position and length. The used scripts were created in-house and will be made available upon reasonable request.

### Calculation of specificity

*Percentage specificity* was calculated by dividing the *percentage reads with indels* of the on-target locus by the sum of *percentage reads with indels* across all three loci (on-target, off-target1, and off-target2). The resulting value answers the question ‘What percentage of the observed indels happen on-target?’, corrected for the total number of reads at each locus.

### DNA cleavage assays

The relative inhibitory effect of the oligo-based inhibitors was determined using *S. pyogenes* Cas9 (NEB), Alt-R CRISPR-Cas9 sgRNA (IDT), EDTA (Merck) and proteinase K (NEB). The substrate was obtained by PCR on CML cells derived gDNA using Q5 DNA polymerase (NEB) and purified on a 10g/L agarose gel using the Wizard SV Gel and PCR Clean-Up system (Promega). All mixtures were made in 1x NEBuffer 3.1 on ice with final concentrations of 80nM Cas9, 320nM sgRNA, 8 nM substrate and 1000nM oligo. For the pre-incubation assays Cas9 and sgRNA were mixed and preincubated for 20 minutes at 25°C and in the meantime the substrate and oligo mixes were made. Equal volumes of RNP and substrate+oligo mixes were combined and incubated at 37°C for 30 minutes. Reactions were stopped by placing them on ice and immediately adding 0.05V of 0.5M EDTA pH=8.0 and 0.05V of proteinase K. Time-point 0 samples were made by immediately stopping the reactions and storing them on ice during the incubation. After stopping samples were incubated at room temperature for 10 minutes and then TriTrack loading dye (ThermoFisher) was added to a final concentration of 1x and 12 μl was loaded on a 1% agarose gel. Gels were run for 45 minutes at 100V in 1xTAE buffer using SYBRSafe (ThermoFisher) as staining agent. Bands were visualized using the UVITEC Alliance (UVITEC) and quantified using the Image Lab software (Bio-Rad) version 6.0.1 build 34. Relative quantity of the remaining substrate was determined by setting the time-point 0 samples of each as 1 and measuring the relative intensity. % digestion was calculated as (1-Irel)*100%. Assays were performed 3 times independently for each substrate.

For the non-preincubated in vitro assay the concentrations of all compounds were the same but the mixtures were substrate + Cas9 and oligo + sgRNA and they were combined on ice before incubation at 37°C.

### Time-shift or time-lapse assay

For the time-lapse assay the same compounds and concentrations were used as for the in vitro assays but the preformed RNP and inhibitor were pre-incubated in 30 second steps before addition of the substrate. Therefore, three pre-mixes were made, Cas9 and sgRNA (RNP), oligo and substrate were diluted in 1x NEBuffer 3.1. The RNP complexes were pre-formed by incubation at 25°C for 20 minutes and 5 μl oligo dilution was put in 1.5 ml tubes (or buffer in the case of the ‘no inhibitor’ control). All samples were put in a 37°C heat block and at different time-points 10 μl of the RNP or 5 μl substrate were added, so all samples had a digestion time of 30 minutes. Reactions were stopped and samples analyzed on gel as described above.

### Electrophoresis mobility shift assay (EMSA)

Final concentrations of the compounds for EMSA were 170nM Cas9, 170nM sgRNA and 500nM oligo. Compounds were added together and pre-incubated at 37°C for 15 minutes after which buffer or the tested compound were added and the samples were incubated at 37°C for 15 minutes. Tritrack loading dye was added and 5 μl was loaded on a 5% polyacrylamide gel in 0.5xTB buffer (45 mM Tris, 45 mM boric acid) and ran at 15mA for 40 minutes. The gel was stained with SYBRGold (ThermoFischer) for 5 minutes, destained in 0.5xTB buffer for 10 minutes and visualized using the UVITEC Alliance (UVITEC).

### Data visualization and statistics

The data presented in this study were visualized using the *datavis.ipynb* Jupyter Notebook file. In addition to *Jupyter Notebook* ^72^ , we used *Python3* ^71^ , *numpy* ^73^ , *pandas* ^74^ , *matplotlib* ^75^ , and *seaborn* ^76^. In cases where results were quantified, triplicates were used for each condition. Where statistics were indicated, those were done by conducting a Shapiro-Wilk test for normality, a Levene’s test for equal variance, a one-way ANOVA, and lastly a Tukey’s range posthoc test. For these statistics, we used *SciPy* ^77^ and *statsmodels* ^78^ in addition to the programs and packages used for the data visualization.

## Supplementary information

**Supplementary Fig. 1.**
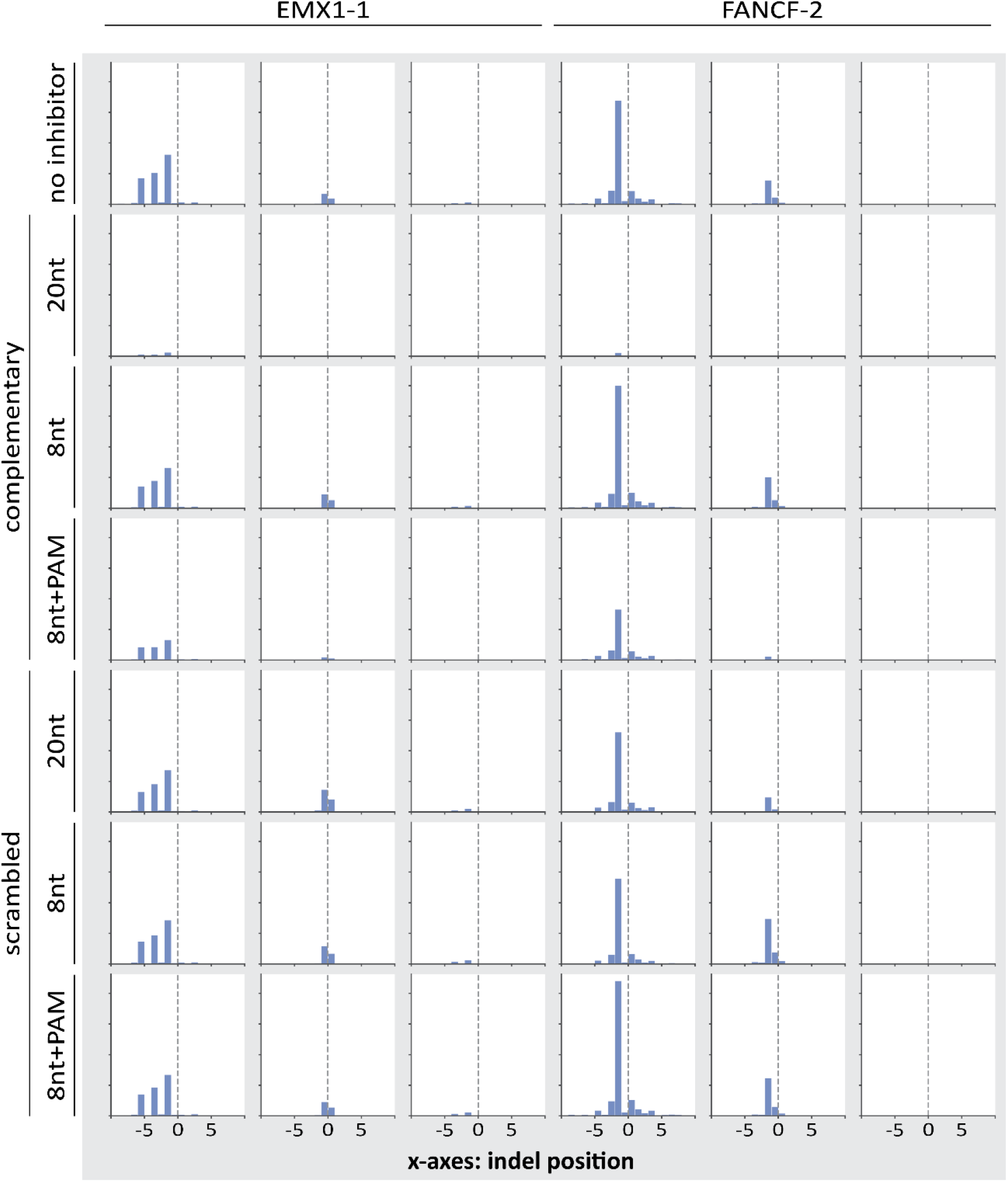
Indel positions distribution. Histograms of the positions of the indels for different DNA oligo-based designs. The ‘0’ position is the Cas9 cut-site for each amplicon. We only displayed indels that occurred within 9 bp from the cut-site. The DNA oligos included in these graphs were delivered at molar concentrations equal to the concentration of Cas9 and guide RNA.

**Supplementary Fig. 2.**
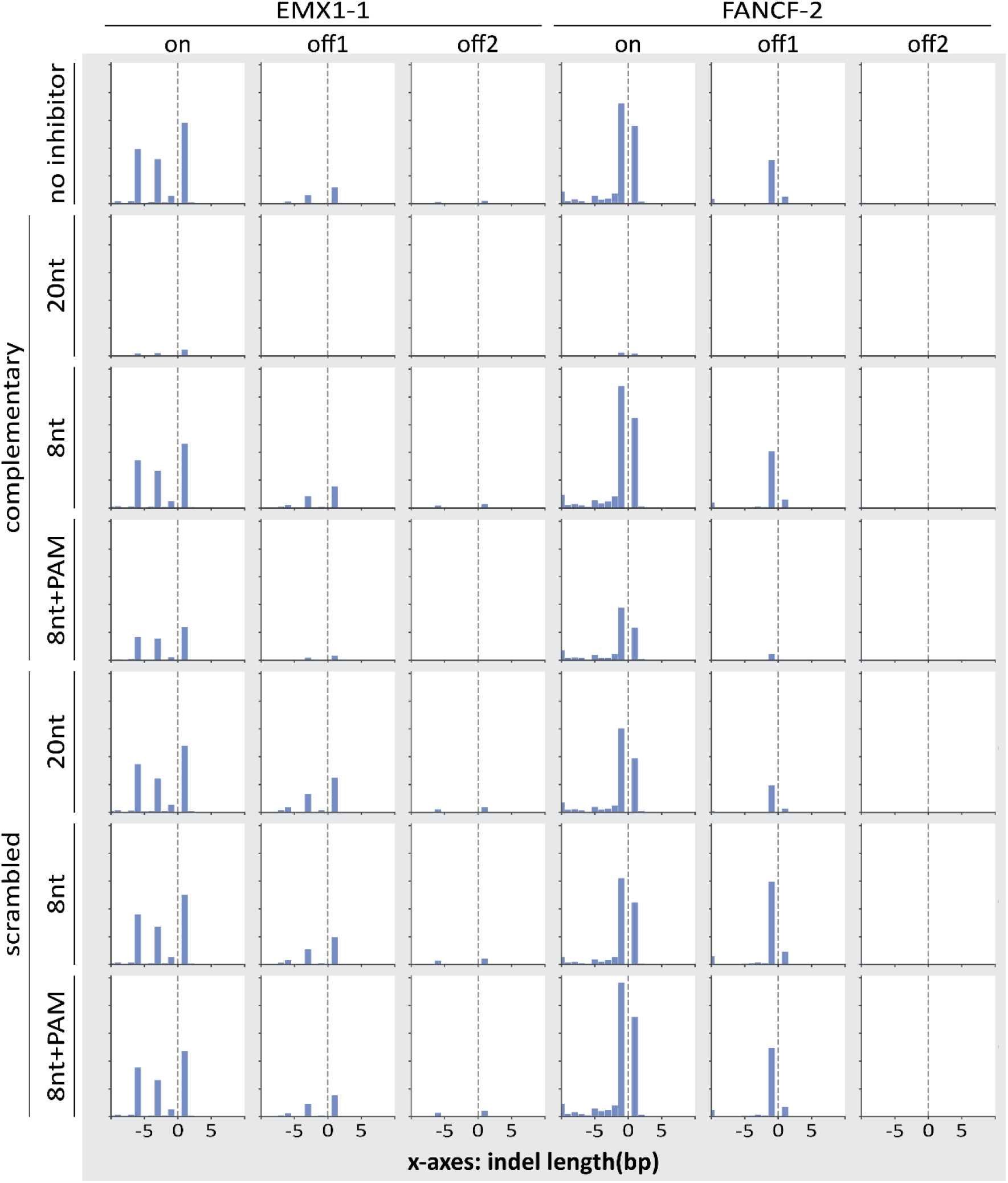
Indel length distribution. Histograms of the lengths of the indels for different DNA oligo-based designs. Negative lengths indicate deletions, whereas positive lengths indicate insertions. A length of ‘0’ would not be an indels, and was not included in these graphs. We only displayed insertions or deletions or 10 bp or shorter. The DNA oligos included in these graphs were delivered at molar concentrations equal to the concentration of Cas9 and guide RNA.

**Supplementary Fig. 3.**
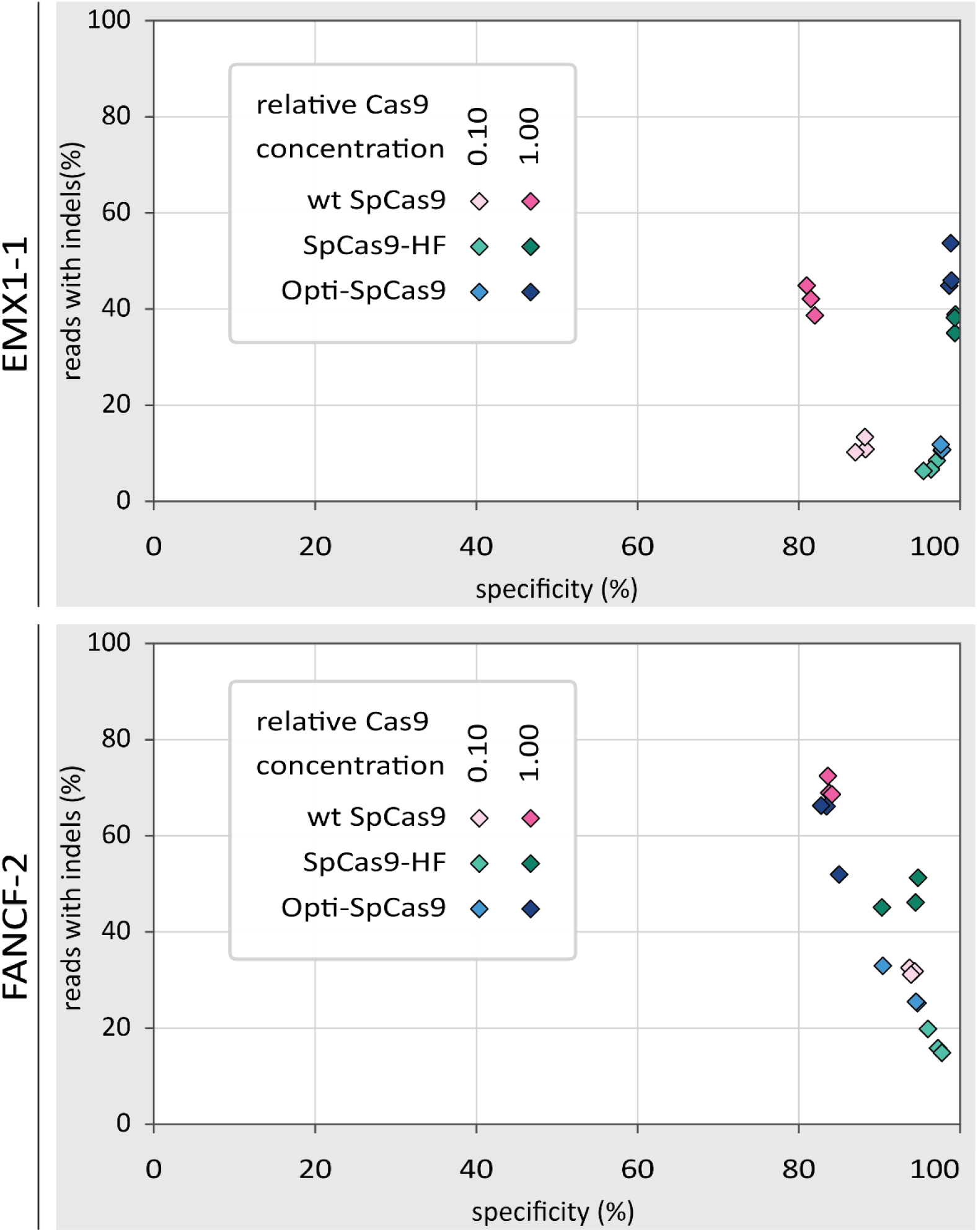
Specificity and activity of engineered Cas9 variants. comparison of *percentage reads with indels* and *percentage specificity* for different Cas9 variants. For each protein, and for each protein concentration, individual replicates are displayed as diamonds with the same color.

**Supplementary Table 1.**
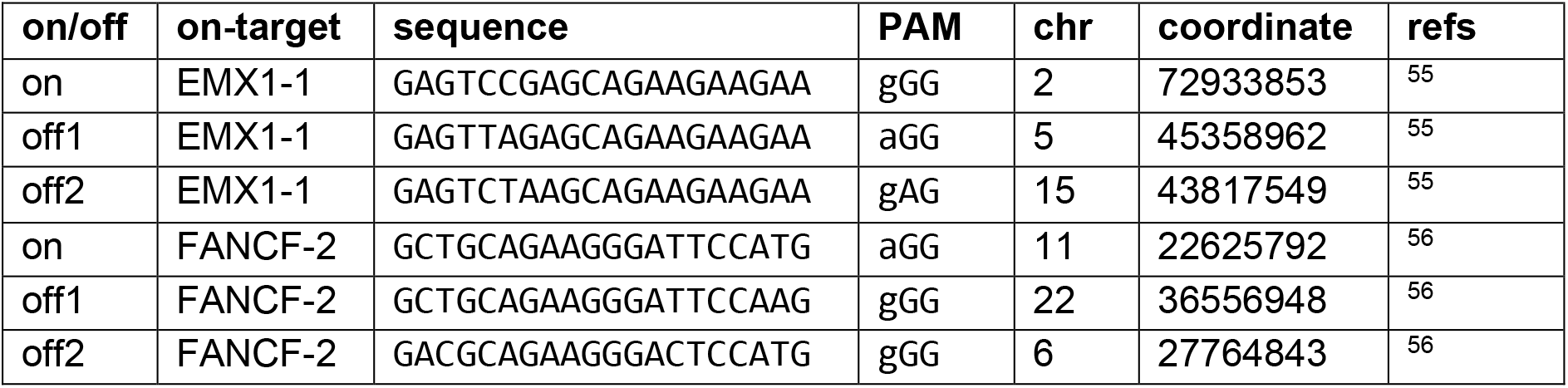
Genomic loci. The genomic loci that were targeted in the *ex vivo* experiment, and which were amplified to provide DNA substrate for the *in vitro* experiments. For each target we list whether it is included as an on- or off-target site, which on-target site it relates to, the nucleotide sequence, the PAM, the chromosome on which it is located, the coordinate of where on the chromosome it is located, and finally references to sources, based on which we decided to include these loci.

**Supplementary Table 2.**
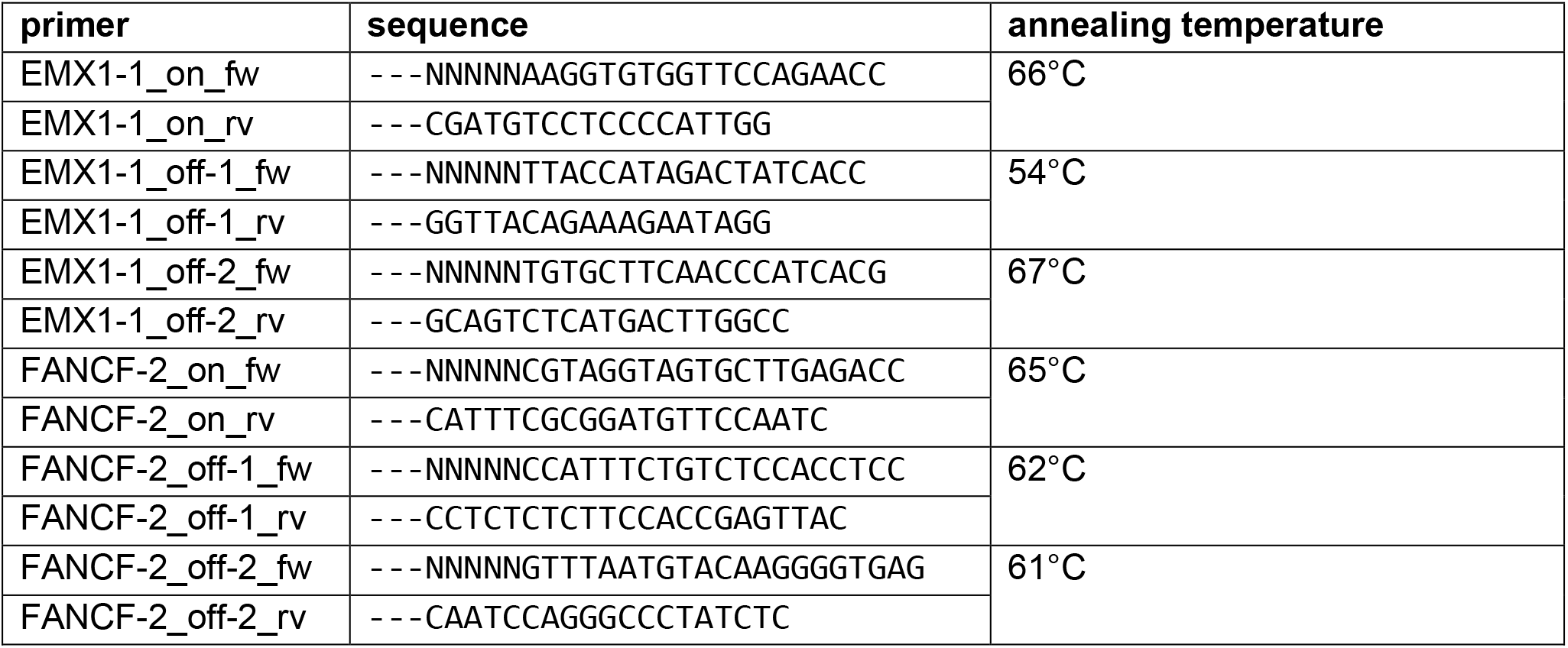
Deep-sequencing preparation primers. The primers that were you to create the amplicons that were sent for deep-sequencing. In the sequences, ‘---’ is used to indicate the upstream sequences that Baseclear B.V. requested for further preparation of the sequencing samples. These sequences are excluded from the amplicon length in the last column. ‘NNNNN’ is used to indicate the 5 nt long barcodes that we used to distinguish between conditions.

**Supplementary table 3.**
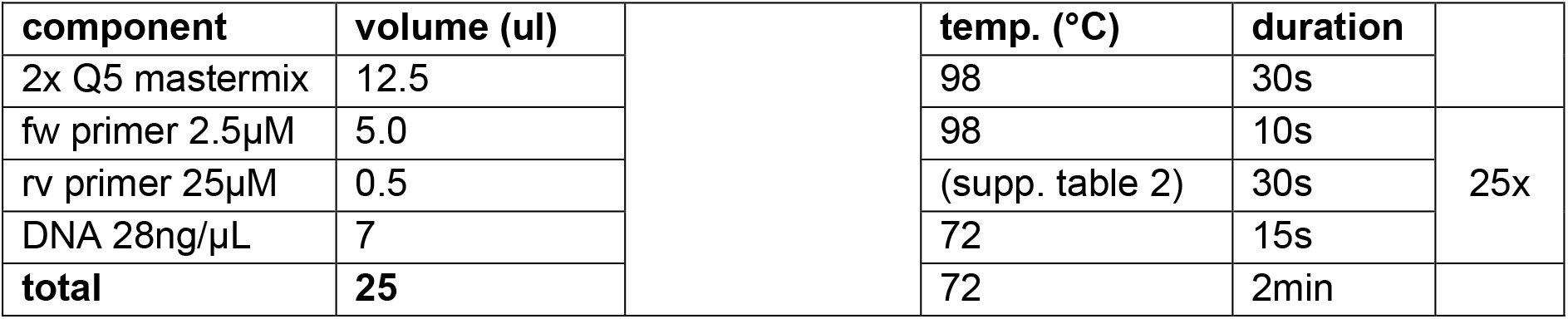
Deep-sequencing preparation PCR conditions. **Left:** The components, their concentrations and used volumes for the 25μL PCR reactions. **Right:** The thermocycler program used for the PCRs.

**Supplementary table 4.**
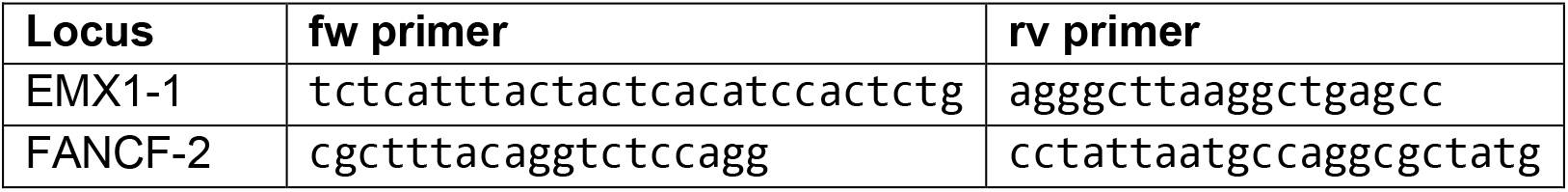
*In vitro* DNA substrate primers. The primers used to amplify the 1500bp DNA substrates from the human genome for the *in vitro* assays. Cleavage by Cas9 results in two fragments of lengths 500bp and 1000bp.

**Supplementary table 5.**
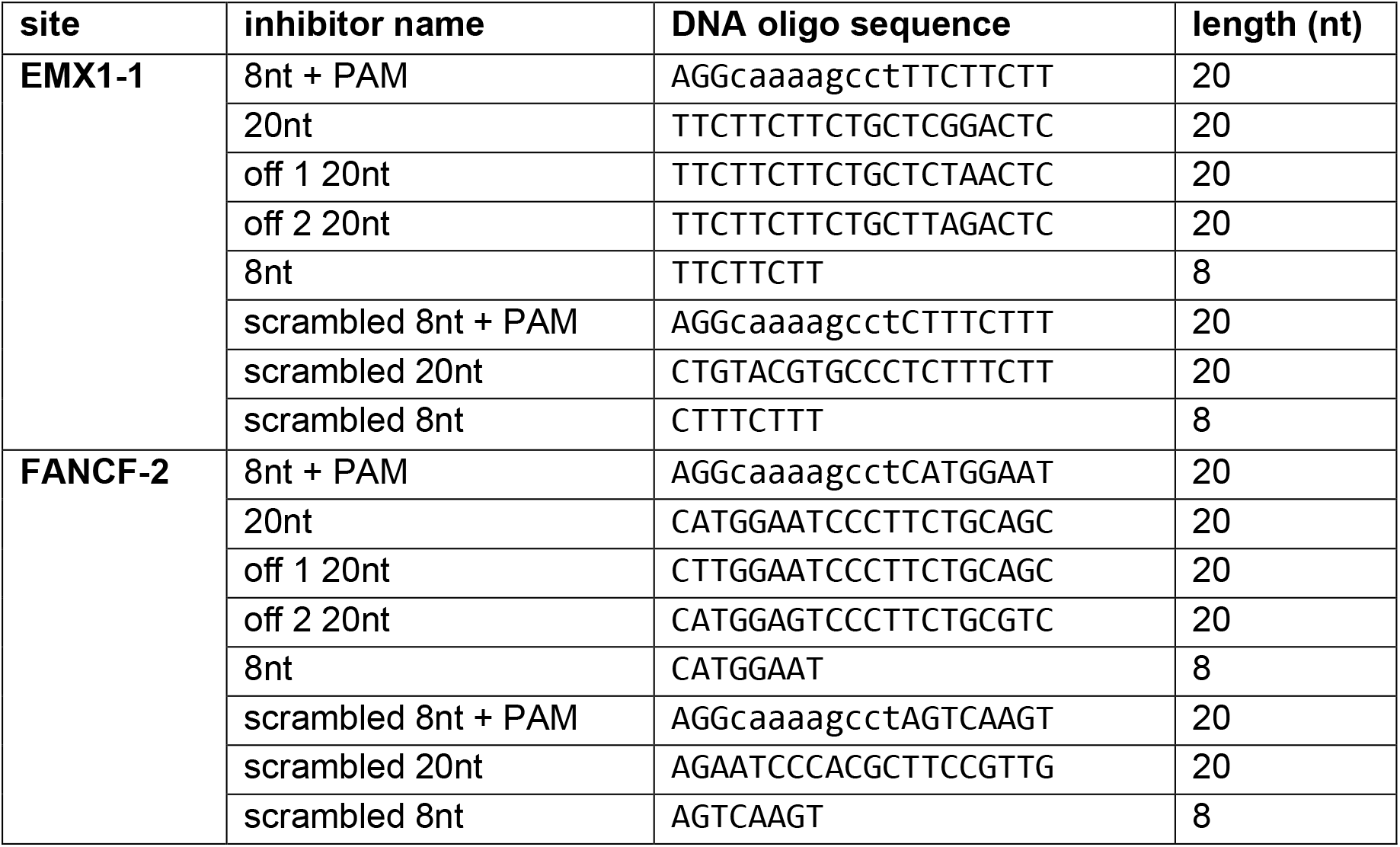
DNA oligo-based inhibitors. The names, sequences, and lengths of the DNA oligonucleotides that were used as inhibitors. In the sequences, both the guide-complementary part and the PAM are in uppercase, but the loop connecting them is lowercase.

